# Analysis of the leaf metabolome in *Arabidopsis thaliana* mutation accumulation lines reveals association of pleiotropy and fitness consequences

**DOI:** 10.1101/2021.06.28.450192

**Authors:** Sydney Kreutzmann, Elizabeth Pompa, Nhan D. Nguyen, Liya Tilahun, Matthew T. Rutter, Mao-Lun Weng, Charles B. Fenster, Carrie F. Olson-Manning

## Abstract

Understanding the mechanisms by which mutations affect fitness and the distribution of mutational effects are central goals in evolutionary biology. Mutation accumulation (MA) lines have long been an important tool for understanding the effect of new mutations on fitness, phenotypic variation, and mutational parameters. However, there is a clear gap in predicting the effect of specific new mutations to their effects on fitness. Here, we complete gene ontology analysis and metabolomics experiments on *Arabidopsis thaliana* MA lines to determine how spontaneous mutations directly affect global metabolic output in lines that have measured fitness consequences. For these analyses, we compared three lines with relative fitness consistently higher than the unmutated progenitor and three lines with lower relative fitness as measured in four different field trials. In a gene ontology analysis, we find that the high fitness lines were significantly enriched in mutations in or near genes with transcription regulator activity. We also find that although they do not have an average difference in the number of mutations, low fitness lines have significantly more metabolic subpathways disrupted than high fitness lines. Taken together, these results suggest that the effect of a new mutation on fitness depends less on the specific metabolic pathways disrupted and more on the pleiotropic effects of those mutations, and that organisms can explore a considerable amount of physiological space with only a few mutations.

**Significance Statement:** As the source of all genetic variation, new mutations are crucial for understanding how organisms adapt to their environment. However, the ways in which new mutations affect the range of metabolic reactions that occur in the cell is unknown. With a combination of gene functional analyses and measurement of the small molecules that drive cellular function and physiology, we find that mutations associated with high fitness are disproportionately found in or near proteins-coding genes that regulate the specific timing and location of gene expression and are less disruptive of cellular physiology.

## Introduction

Adaptation requires mutations that both beneficially affect a certain trait under selection and, importantly, do not have large deleterious effects on other traits. Mutations in or near proteins that regulate the timing and location of gene expression may satisfy both of these conditions. Transcription regulatory proteins and trans-acting DNA elements interact with the cis-regulatory DNA to direct the spatial and temporal gene expression in eukaryotes. This expression occurs in a highly specific manner as the DNA regulatory elements of genes and trans-acting regulatory proteins act in a combinatorial fashion. A growing number of studies have found that mutations that affect the expression of trans-acting loci (Streisfeld, Liu, and Rausher 2011; Streisfeld and Rausher 2009; Whittall et al. 2006; Quattrocchio et al. 1999; Hoballah et al. 2007) and cis-regulatory DNA (Des Marais and Rausher 2010; Hopkins and Rausher 2011) make important contributions to adaptive evolution.

In the case of transcription regulatory proteins, while some are widely expressed, a majority are expressed in only a few tissues or under specific environmental conditions (Streisfeld, Liu, and Rausher 2011). Likewise, cis-regulatory structure is highly module and mutations that occur in the cis-regulatory regions of a gene generally affect only the the expression of other modules (Wittkopp and Kalay 2011). Due to this tissue-specific functioning, mutations that are in or near genes that affect the expression of most transcription regulatory proteins are less likely to have widespread pleiotropic consequences and are thus less likely to be deleterious (Wagner and Lynch 2008). Given the demonstrated importance of regulatory evolution for adaptation, we would like to understand the prevalence of mutations that affect gene regulation in the mutational spectrum (i.e., the range of spontaneous mutations that occur) and to predict the effect of mutations on organismal physiology and fitness.

An important tool for examining the types and effects of new spontaneous mutations are mutation accumulation (MA) lines (Halligan and Keightley 2009). MA lines are created when sublines of an original progenitor are allowed to accumulate spontaneous mutations in the absence of selection through many generations. These lines have shed light on the distribution of phenotypic effects (Chang and Shaw 2003; Shaw, Geyer, and Shaw 2002; Weng et al. 2021), the mutational spectrum (Weng et al. 2019; Schrider et al. 2013; Denver et al. 2006; Grey Monroe et al. 2020; D. T. Nguyen et al. 2020), and the average fitness effect of new mutations (Rutter, Roles, and Fenster 2018; Rutter et al. 2012; Roles et al. 2016; Rutter, Shaw, and Fenster 2010) (reviewed in (Katju and Bergthorsson 2019; Eyre-Walker and Keightley 2007)).

The *Arabidopsis thaliana* MA lines (Chang and Shaw 2003; Shaw, Geyer, and Shaw 2002) offer an opportunity to connect mutation to fitness through physiology. The *A. thaliana* MA lines have been fully sequenced (Weng et al. 2019) and the fitness measured in field trials (Rutter, Roles, and Fenster 2018; Rutter et al. 2012). However, there has been little use of this resource to examine the mutational effect of fitness through physiology. In accordance with theory (Fisher 1930) and supported by empirical studies (e.g. (Cooper, Nickerson, and Eichler 2007)), we predict that lines that have higher fitness than the progenitor should also have proportionally more mutations that display low levels of pleiotropy, and these mutations may be enriched in genes responsible for regulation. With these high and low fitness lines and their associated mutations, we ask 1) are mutations in or near regulatory proteins enriched in MA lines that have high relative fitness compared to the unmutated progenitor? and 2) Are there more overall changes in metabolic output in the low fitness lines compared to the high fitness lines?

In an analysis of the gene ontology of the mutations in or near the genes in the high and low fitness lines, we find that mutations in MA lines with high relative fitness are enriched in proteins with transcription regulator activity. We also find an intriguing (but possibly underpowered) trend where high fitness lines have fewer changes to their metabolic pathways than the low fitness lines, suggesting that the mutations in high fitness lines are less pleiotropic than mutations in the low fitness lines. Contrary to our initial assumption, we find no clear connection between the mutations and their expected effect on metabolism. Taken together, these results suggest that mutations that affect transcription regulator activity are common in the mutational spectrum and that lines that harbor these mutations have less disrupted metabolic expression. Although we find mutations in or near highly annotated genes, we see no predictable effect on metabolism, suggesting our annotations of gene function are incomplete.

## Results and Discussion

The ways in which mutations affect organismal fitness is of great interest in evolutionary biology and molecular evolution. Although other evolutionary forces determine the fate of the mutations after they arise, bias in the mutational spectrum can affect the composition of the genome (D. T. Nguyen et al. 2020), how the composition of the genome affects mutation rate (Grey Monroe et al. 2020), and polymorphism in populations (Weng et al. 2019). In our analyses, we seek to connect the composition of spontaneous mutations in the *A. thaliana* MA lines with high and low relative fitness (measured under field conditions) and the effect of those mutations on organismal metabolism.

### High fitness lines are enriched in transcription regulator activity

In a Gene Ontology (GO) term analysis for molecular function (Mi et al. 2021) we found genes in the high fitness lines submitted for metabolomics in this study have a 7.7-fold enrichment (Fisher’s Exact Test with Bonferroni correction, p<0.01) in transcription regulatory activity (8 of the 16 mapped gene IDs and represent between 25-50% of the mutations in each line). We repeated this GO term analysis for all 18 high fitness MA lines (Supplementary Table 2) and found that together they also have a greater than 3.1-fold enrichment for transcription regulator activity (p<0.001, 21 of 114 mapped gene IDs have transcription regulator activity). We identified no other significant enrichment in the genes in the high fitness lines. The same analyses on all 11 low fitness lines revealed no enrichment in any activity and only one out of 32 genes with mutations in the low fitness lines has transcriptional regulatory activity.

These results suggest that mutations in or near genes encoding transcriptional regulatory proteins make up a substantial portion of the mutational spectrum in MA lines with high relative fitness and are less prevalent in lines with low fitness. Proteins with transcriptional regulatory activity assemble into varied complexes consisting of other proteins and cis-regulatory sequences in a combinatorial fashion to tightly control the timing and location of gene expression (Brkljacic and Grotewold 2017). In this way, a small handful of transcription factors can control the expression of a wide variety of genes. The vast majority of the mutations in our mutation accumulation lines are intergenic, and thus these mutations likely affect the regulation of the identified proteins with transcription regulatory activity (the mutations are in the cis-regulatory regions or in trans enhancers or repressors. The combinatorial nature of these regulatory networks have led many to hypothesize that mutations that affect gene regulation could play an outsized role in adaptive evolution (Jeong et al. 2008; Wray et al. 2003). The evolutionary genetics literature has many examples implicating mutations in proteins with transcriptional regulatory activity in adaptations, including flower color evolution (Streisfeld, Liu, and Rausher 2011; Wessinger and Rausher 2013; Lin and Rausher 2021; Quattrocchio et al. 1999; Whittall et al. 2006), adaptation to high temperature (Koini et al. 2009), cold acclimation (Van Buskirk and Thomashow 2006), drought (Haake et al. 2002; Leng and Zhao 2020; Jan et al. 2019), Zinc deficiency (Inaba et al. 2015), the interface between stress and growth response (Danisman 2016) and many more. Given the purported role of proteins with transcriptional regulatory activity (Wagner and Lynch 2008), and their prevalence in the mutational spectrum in lines with high relative fitness, these potentially low pleiotropy mutations are readily available for adaptation.

### Metabolic disruption in high and low fitness lines

We find a consistent trend of more metabolic subpathways disruption in the low fitness lines than in the high fitness lines. The low fitness lines have an average of 25 disrupted subpathways compared to an average of 15.3 disrupted subpathways in the high fitness lines when we consider all metabolites that approach significance (two-sided unpaired t-test, p-value = 0.02) (Table 1, Supplementary Table 3). This pattern of a higher average number of disrupted metabolites in low fitness lines is consistent at the super pathway, subpathway, and individual metabolite levels, regardless of whether we include only metabolites that reach a the typical significance threshold p≤0.05 or metabolites that approached significance (0.05<p<0.10 (Table 1, Supplementary Table 4). Overall, these results suggest that mutations in the low fitness lines affect more parts of metabolism than mutations in high fitness lines.

**Table 1:** The Number of enriched metabolic subpathways and individual metabolies calculated for metabolites that met the 0.05 signifinace threshold and those that met the 0.1 threshold. Significant difference between the number of enriched subpathways that reach the 0.1 threshold denoted with * (p<0.05 unparied t-test).

| Line number | Relative Fitness | Enriched Subpathways $p < 0.05$ | Enriched Subpathways $p < 0.1$ * | Total metabolites $p < 0.05$ | Total metabolites $p < 0.1$ |
| --- | --- | --- | --- | --- | --- |
| 44 | Low | 10 | 21 | 13 | 37 |
| 73 |  | 20 | 29 | 32 | 60 |
| 75 |  | 12 | 25 | 13 | 39 |
| 53 | High | 14 | 17 | 16 | 25 |
| 61 |  | 10 | 14 | 18 | 35 |
| 119 |  | 7 | 15 | 9 | 25 |

The precise set of regulatory proteins is controlled in a tissue-specific manner and the level of constraint in different regulatory elements is reliably predicted by level of integration into gene regulatory networks (Streisfeld, Liu, and Rausher 2011). Taken together, the lower levels of pleiotropy and enrichment in mutations in or near genes with transcription factor activity in the high fitness lines is consistent with studies that find transcription factors are over-represented in adaptive evolution due to their low-pleiotropic nature (Smith 2016; Wagner and Lynch 2008; Cheatle Jarvela and Hinman 2015).

### Pleiotropic effects of mutations on metabolic pathways

While the high and low fitness lines have no average difference in the number of unique, non-transposable element mutations (average of 9.0 for high fitness and 10.7 for low fitness lines, Supplementary Table 1), there is considerable variance in the number of mutations among the lines. There is no relationship between the number of mutations and the number significantly disrupted metabolic subpathways (F-statistic 0.017, p = 0.9026). On average, each mutation affects multiple metabolic pathways. Mutations in four of the six lines affect 1.5-3.5 metabolic subpathways per mutation (Supplementary Table 1). High fitness line 119 and low fitness line 44 both affect fewer than one metabolic subpathways per mutation. With this small sample size and the confounding effects of individual mutations in each line, we cannot know with certainty how many pathways each mutation affects. However, based on these data, it appears that *A. thaliana* can explore a considerable amount of physiological space with only a few mutations.

### Spontaneous mutations cause excess down-regulation of metabolic pathways

We found that overall, mutations caused significantly more down-regulation to metabolic pathways than up-regulation in both the high and low fitness lines (Figure 1, blue indicates down-regulation, red is up-regulation, Chi-square p<0.007). Down-regulated metabolites are 4-times more common than up-regulated metabolites and there was no significant difference in the number of up- and down-regulation in a comparison of the high and low fitness lines. It is possible that mutations are more likely to lead to down-regulation because mutations tend to disrupt steps in metabolic pathways.

**Figure 1:**
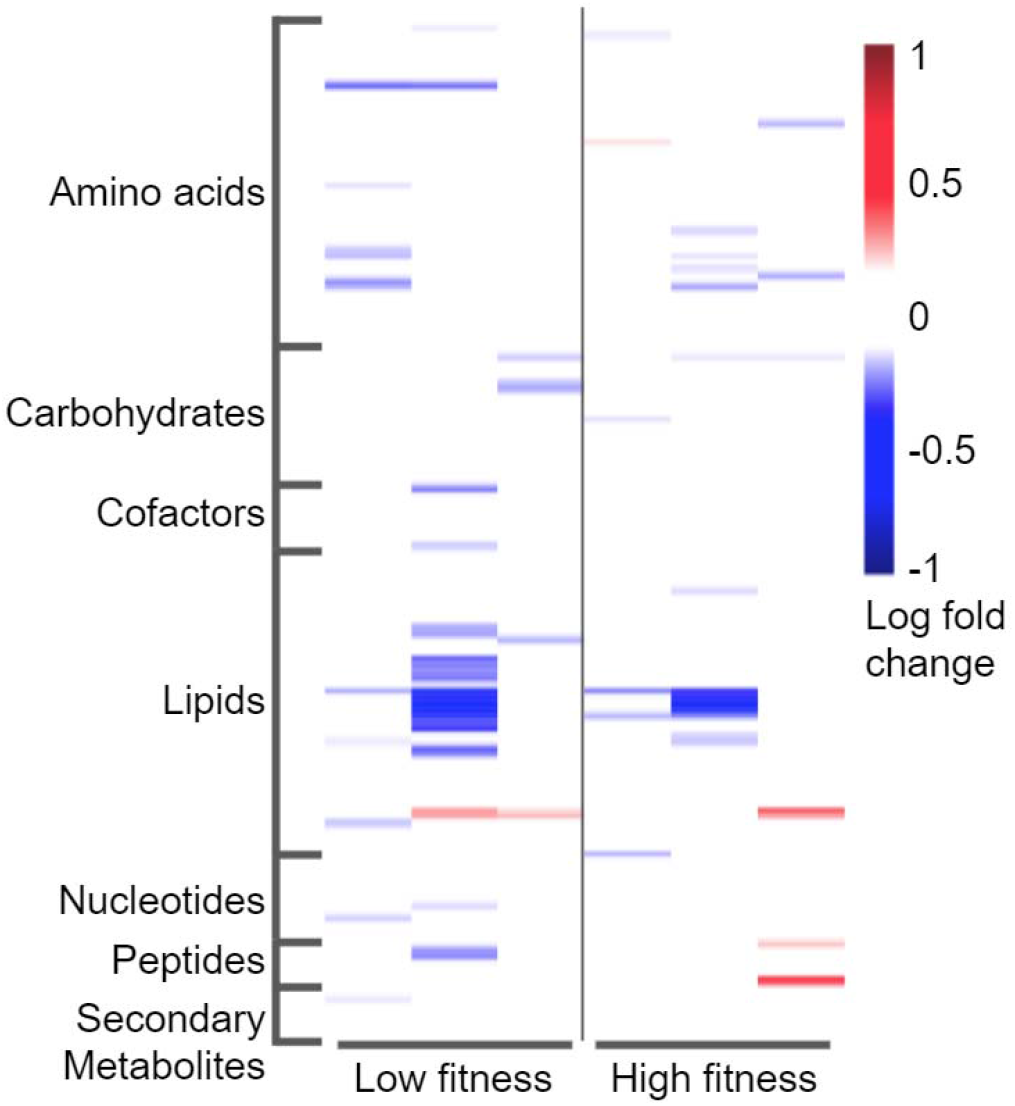
Log fold change of all significant and near metabolies. The columns from the left to right are the low fitness lines (44, 73, 75) and high fitness lines (53, 61, 119). The rows are ordered based on superpathway (full data in Supplemental Table 3).

While it might be reasonable to assume that general degradation of metabolic function is deleterious, there are many instances of loss of function alleles contributing to adaptation (Stower 2013; Xu et al. 2019; Grey Monroe et al. 2020; Monroe et al. 2018). In *A. thaliana*, loss-of-function alleles are common in coding genes, and an estimated 1% of loss-of-function alleles are under positive selection (Xu et al. 2019). Therefore, without further measure of allele frequency changes in the MA lines following selection, it is not possible to speculate on the evolutionary importance in an overall reduction in metabolic activity.

### Spontaneous mutations affect metabolism in varied ways

The effect of new spontaneous mutations on metabolism is varied (Fig. 1 and 2) and suggests that the mutations accumulated in these lines are idiosyncratic with respect to effect on metabolic pathways. In a comparison of the six lines with PCA analysis (Figure 2) the high and low fitness lines are intermixed suggesting that there are not consistent changes to the types of metabolic disruption that led to high or low fitness. These results are also consistent with the heatmap generated from the log-fold data for all metabolites (Supplementary Figure 1). These results suggest that the spontaneous mutations were indeed random with respect to their effect on metabolism.

**Figure 2:**
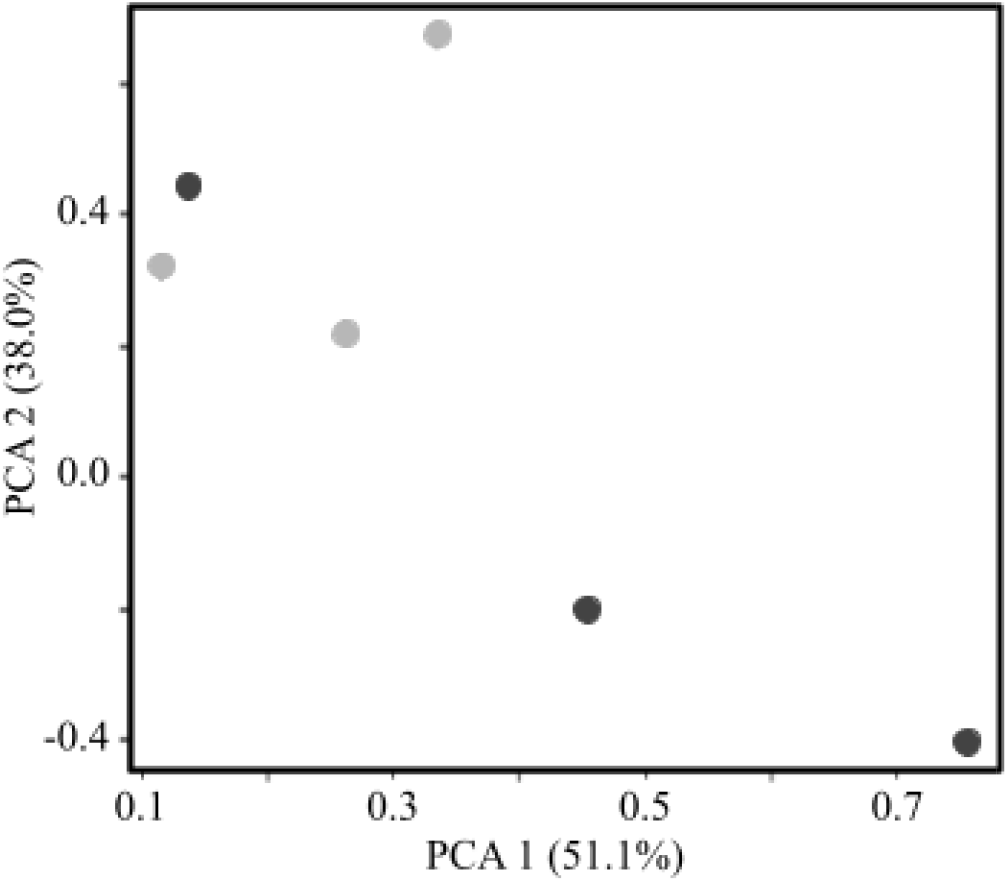
Principal component analysis (PCA) of the fold-change metabolomics data with the percentage of variance explained by each principal component is denoted in the parentheses. The lines with high relative fitness (dark grey points) and the lines with low fitness (light grey points) do not group according to fitness.

**Figure 3:**
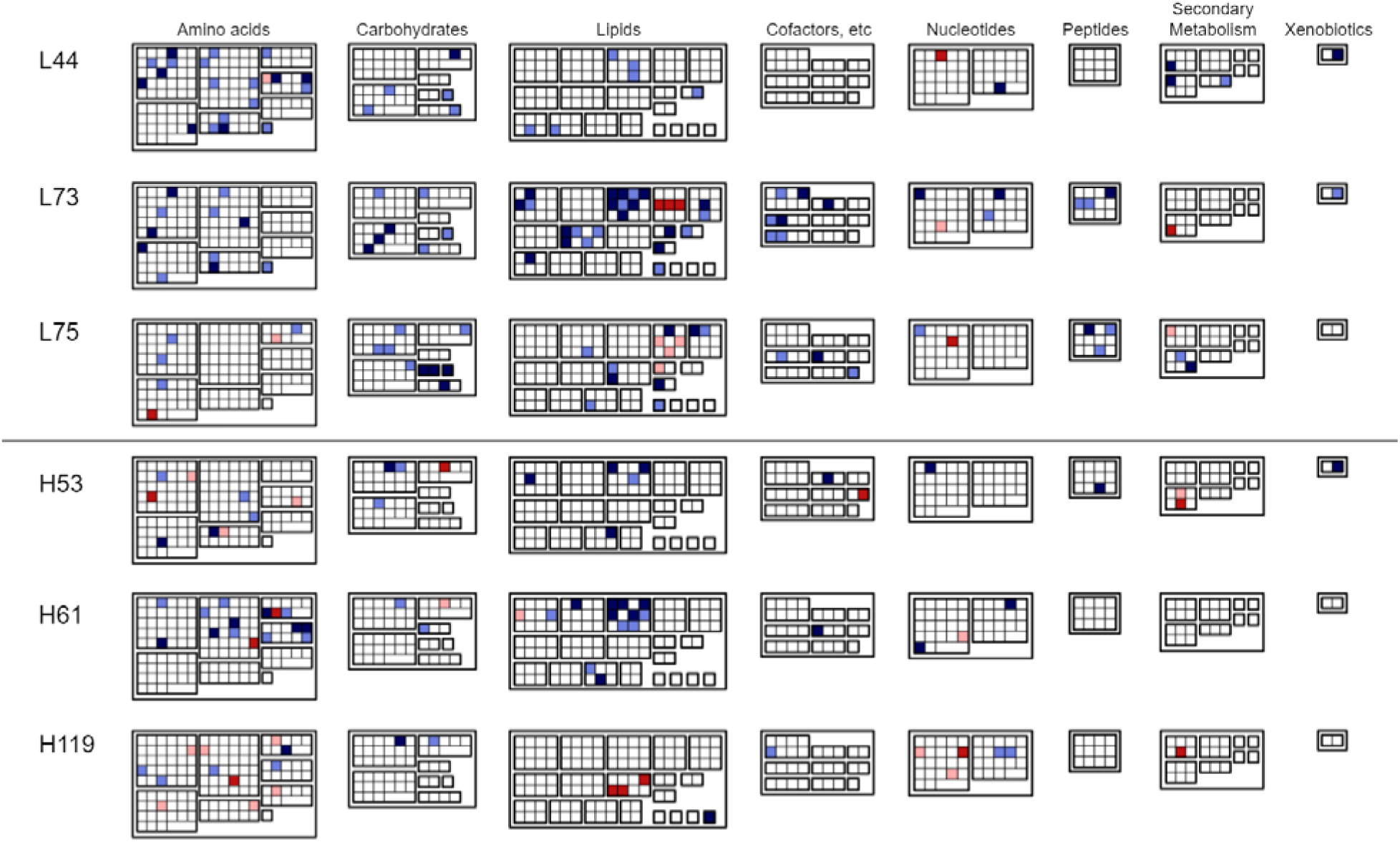
Up- (reds) and down-regulation (blues) metabolites in their super pathways context with subpathways in individual boxes. Lighter colors are metabolites that approach significance. We excluded super pathways from this figure that did not show any significant or near-significant differences (hormone metabolism and partially characterized molecules metabolism). A key of each metabolite is found in Supplementary Figure 2.

### The potential and limitations of metabolomics studies in understanding the effects of spontaneous mutations

Understanding and predicting the effect of mutations is a primary goal across many fields of biology (Eyre-Walker and Keightley 2007). In this study, we used metabolomics to try to close the gap between a mutation and its effect on fitness. We find that although our results are consistent across all metrics, our study is underpowered and we are basing our connections between metabolic disruption and fitness based primarily on consistent trends in the number of disruptions (and not on strong statistical or large fold-change effects) in the comparison of the lines with high relative fitness and low relative fitness. This lack of power is, in retrospect, unsurprising given each mutation accumulation line has only a small number of mostly intergenic mutations. We expected to find larger effects given the consistent trends in relative fitness observed across environments in these lines (Supplemental Table 1). Nonetheless, we believe these results are a valuable step toward integrating metabolomics experiments into evolutionary analyses. The cost of metabolomics experiments still limits the number of biological replicates and developmental time points that can be measured for a large number of samples. As prices continue to decline, this methodology is poised to provide hundreds of physiological measures that can link to the abundance of sequencing data now available.

Even with hundreds of metabolites measured in this study in well-known pathways and the well-annotated *A. thaliana* genome (Lamesch et al. 2012) we failed to infer any direct connections between mutations in or near enzymes with an effect on metabolism (full list of AT TAIR numbers and functional annotation available in Supplementary Table 2). Among the reasons for this failed connection could be other sources of variation (Becker et al. 2011; Davies et al. 2016), that most of the mutations in the MA lines in this study are intergenic and have direct or indirect effects on the expression genes far away, that the small number of mutations failed to disrupt any known pathways or pathways measured in this study, or that our annotations of gene function are woefully incomplete.

This study only measures the metabolic effects in leaf tissue at one time point on only six of the 107 *A. thaliana* MA lines, so the conclusions relating to the pleiotropic nature of the mutations should be viewed in that light. We also chose lines that showed consistent relative fitness in field trials (Rutter, Roles, and Fenster 2018); however, most of the lines had strong genotype-by-environment effects. Additionally, studies on the effect of new, spontaneous mutations on transcriptional expression have shed light on some of the biases of the mutational spectrum (Zalts and Yanai 2017; Huang et al. 2016), but lack fitness measures to connect genotypic changes to fitness effects. Future work that includes transcriptional studies combining metabolic effects with sampling of additional tissues and an analysis of more lines under many conditions will further elucidate the effect of spontaneous mutations on gene expression, fitness, and physiology.

## Materials and Methods

### Choice in plants to grow and analyze

Of the 107 total *A. thaliana* mutation accumulation (MA) lines (Chang and Shaw 2003; Shaw, Geyer, and Shaw 2002), we chose lines that showed relative fitness that was higher and lower in most environments compared to the unmutated Col-0 progenitor (Rutter et al. 2018) for metabolic analysis (Supplementary Table 1). The focal MA lines in this study (line numbers 44, 53, 61, 73, 75, and 119) are numbered based on the original propagation line numbers (Chang and Shaw 2003; Shaw, Geyer, and Shaw 2002). Briefly, in a previous study, Rutter et al. (2018), all MA lines were grown in four temporal environments at a single field site and plant fitness of the MA lines was compared to the unmutated progenitor. While most lines displayed genotype-by-environment dependent fitness, for this study we chose lines that consistently ranked among the highest or lowest in relative fitness compared to the progenitor (Supplementary Table 1). As defined in this manuscript, the “high fitness lines” (numbers 53, 61, and 119) had higher relative fitness than the progenitor in at least three of four environments and the “low fitness lines” (lines 44, 73, and 75) had lower relative fitness than the progenitor in at least three of four environments (Rutter et al. 2018). These lines have accumulated an average of 17 mutations over 30 generations of mutation accumulation (Weng et al. 2019).

### GO Term Enrichment comparison in high and low fitness lines

To identify gene ontology terms that might be over- or under-represented in the high or low fitness lines, we entered the *A. thaliana* gene model names for mutations (or the gene model names surrounding intergenic mutations) into the Protein Analysis Through Evolutionary Relationships (PANTHER) resource (Mi et al. 2021) (gene names found in Supplementary Table 2). We compared the set of mutations in the high and low fitness for biological processes, molecular function complete, and cellular component complete with the Fisher’s exact test with Bonferroni correction for multiple testing. The analyses were performed separately on the set of three high and three low fitness lines of focus in this study. We also analyzed the additional 15 lines that had average relative fitness that was higher and 8 with relative fitness lower than the progenitor (Rutter, Roles, and Fenster 2018). Here we report the only significant enrichment result we identified.

### Plant growth and tissue harvest

Individual seeds from the 25th generation of mutation accumulation were sown on moistened soil (Promix BX with Osmocote fertilizer added per manufacturer’s instructions), randomized in a 24-cell pallet and allowed to imbibe. The seeds were placed in the dark for 3 days at 4 C to stratify and overcome dormancy. Plants were grown for 3 weeks under long day conditions (16h light) at 18 C. The plant rosettes at time of harvest were vigorous and did not display any obvious signs of stress or growth defects. To control for the developmental stage, all above-ground tissue was harvested before the plant began bolting and was immediately frozen on liquid nitrogen. Each of the high and low fitness lines had four biological replicates and the progenitor had six. The tissue was stored at −80 C until further processing. Frozen leaves were pulverized on liquid nitrogen and added to a frozen collection tube and stored at −80 C until shipment. Samples were shipped on dry ice to Metabolon Inc. (North Carolina). Frozen samples were lyophilized and each sample was standardized with dry weight.

### Metabolon sample extraction and metabolite identification

Briefly, Metabolon Inc. extracted the samples with the MicroLab STAR System (Hamilton Company) with the internal standards. Each extract was divided into five fractions: two were analyzed using acidic positive ion conditions (one optimized for hydrophilic compounds, one for hydrophobic), two used basic negative ion conditions (one optimized for hydrophilic compounds, one for hydrophobic), and one used for a backup. Metabolomic analyses were performed using Ultra High Performance Liquid Chromatography-Tandem Mass Spectroscopy (UPLC MS/MS). These methods utilized a Waters Aquity UPLC and Q-Exactive high resolution mass spectrometer (Thermo Scientific) with a heated electrospray ionization (HESI-II) source. The MS analysis used dynamic exclusion, alternating between MS and data-dependent MS^n^, and varied between 70-1000 m/z. The MS/MS scores were derived from a comparison of the ions in the experimental samples and the ions in a known library spectrum (within the Metabolon LIMS system. The MS/MS scores (forward and reverse), retention time/index, and molecular mass (m/z) were all processed using Metabolon’s software and compared to purified standards within the internal library to identify each metabolite. Peaks were quantified by integrating the area-under-the-curve. Samples analyzed over multiple days were corrected in run-day blocks and normalization based on internal standards.

### Identification of disrupted metabolites and pathways

A total of 386 known compounds were identified in this dataset. Missing data points were given the minimum observed value for each compound and all data were log transformed. Each metabolite was categorized into one of nine super pathways (e.g. Amino Acid, Carbohydrate, Peptide, Secondary Metabolite) and into one of 58 subpathways (e.g. Serine family amino acid, TCA cycle, Dipeptide, Benzenoids) (Supplementary Table 3) (Kanehisa and Goto 2000). Biological replicate measures of each metabolite were compared to the value in the progenitor with Welch’s two-sample t-test. A metabolite was judged to be disrupted in a certain line if its value was significantly different from the value in the progenitor based on the t-test based on an alpha value of 0.05.

We took two approaches to identify enrichment in metabolic subpathway disruption to balance Type I and Type II errors. With a pathway enrichment metric (Xia and Wishart 2011), the number of significant (p≤0.05) compounds were calculated for individual metabolic subpathways. The Enrichment score (Xia and Wishart 2011) compares the ratio of significant metabolites in a pathway (k) to total detected metabolites in that pathway (m), and standardized it by considering the total number of significant metabolites in the entire dataset (n) and the total metabolites detected in the dataset (N) (Eq 1) (results averages in Table 1 and calculations in Supplementary Table 4).

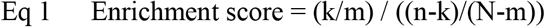

Due to the complex nature of biological interactions, and the small sample size used in this study, the traditional enrichment score is likely conservative (T.-M. Nguyen et al. 2019). The traditional enrichment score ignores individual pathways with many metabolites that approach significance in the Welch’s two-sample t-test (p<0.1). As a less conservative test, we also calculated an expanded enrichment score that includes metabolites that approach significance (Table 1, Supplementary Table 4). This test allowed us to identify metabolic subpathways that have sets of metabolites that are disrupted and share common biological function. The trend that we observe that lines with relatively low fitness have more disruption is consistent across the traditional enrichment analysis, the extended analysis, and the number of metabolites.

### Comparison of up and down regulation of metabolic pathways

We compared the number of metabolites that were significantly up- or down-regulated as compared to the Col-0 progenitor with a Chi-square statistic. The null expectation is that spontaneous mutations will cause an equal number of up- and down-regulation in metabolites.

## Data Accessibility Statement

All data used in this study (including raw metabolic measures) is represented in the supplementary tables. The datasets will also be uploaded to the Dryad repository upon acceptance.

## Author Contributions section

All authors contributed to the writing, discussion, and revisions of this manuscript.

Author C.F.O.-M., M-L.W. and C.B.F. contributed to experimental design and took part in every contribution listed below.

Authors S.K., E.P, N.D.N., and L.T. contributed to data collection and analysis.

M.T.R. contributed to data analysis.

## Acknowledgments

We thank Detlef Weigel for thoughtful comments and for the generation of the sequencing data. This work was supported by the National Science Foundation (DEB 2017485 to C.F.O.-M, OIA EPSCoR 1920954, DEB 1257902 to C.B.F., DEB 0844820 to C.B.F., DEB 1258053 to M.T.R., and DEB 0845413 to M.T.R.); and the National Institute of Health (INBRE 2P20GM103443-19).

