## Supplemental Figures 1 and 2 for "Analysis of the leaf metabolome in *Arabidopsis thaliana* mutation accumulation lines reveals association of pleiotropy and fitness consequences"

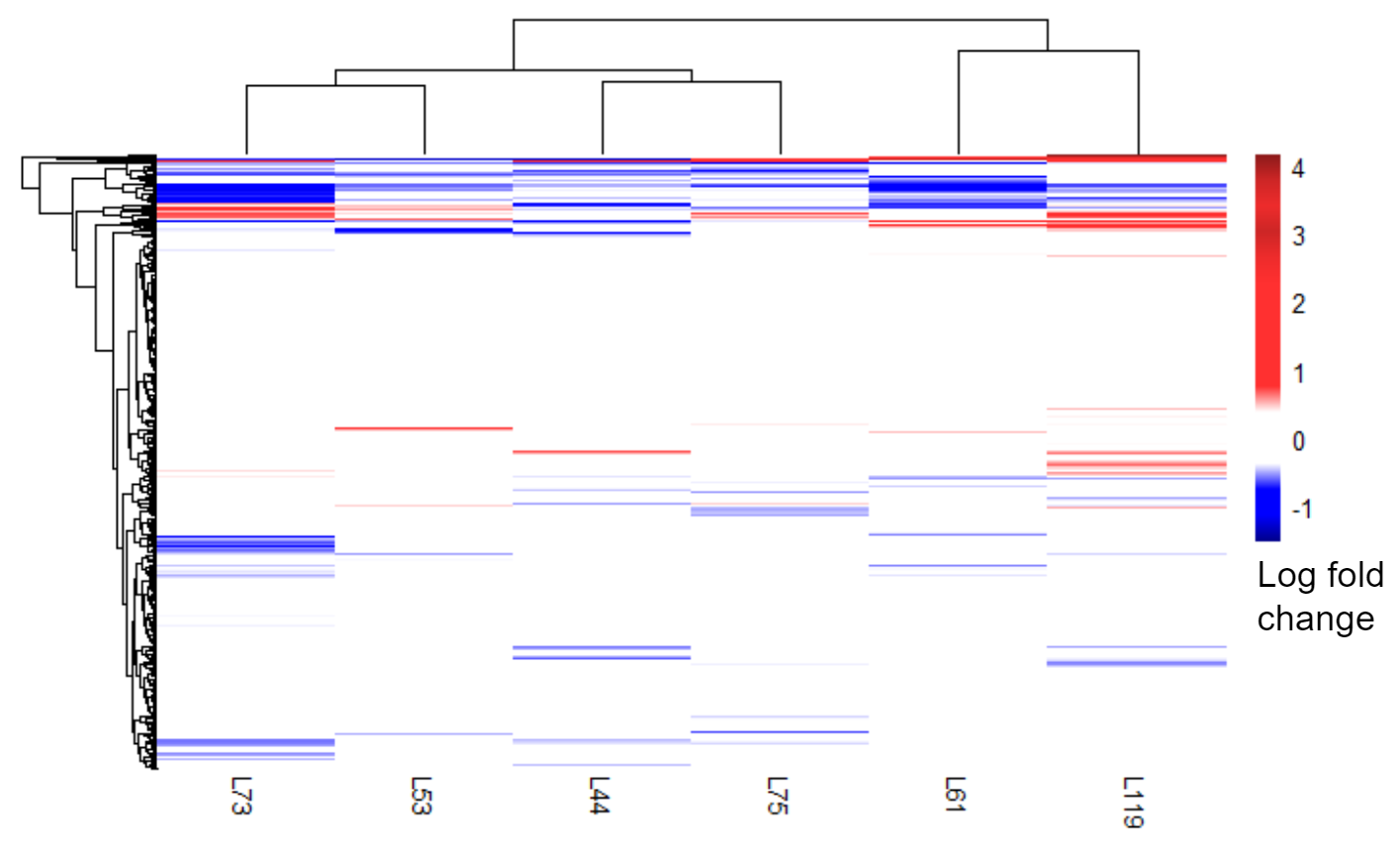

Supplementary Figure 1: Log fold change of all metabolites. The columns are labeled based on the MA line number and are based on Euclidean distance. The rows represent metabolites and are also clustered based on Euclidean distance. The high fitness (L53, L61, L119) and the low fitness lines (L44, L73, L75) do not cluster together

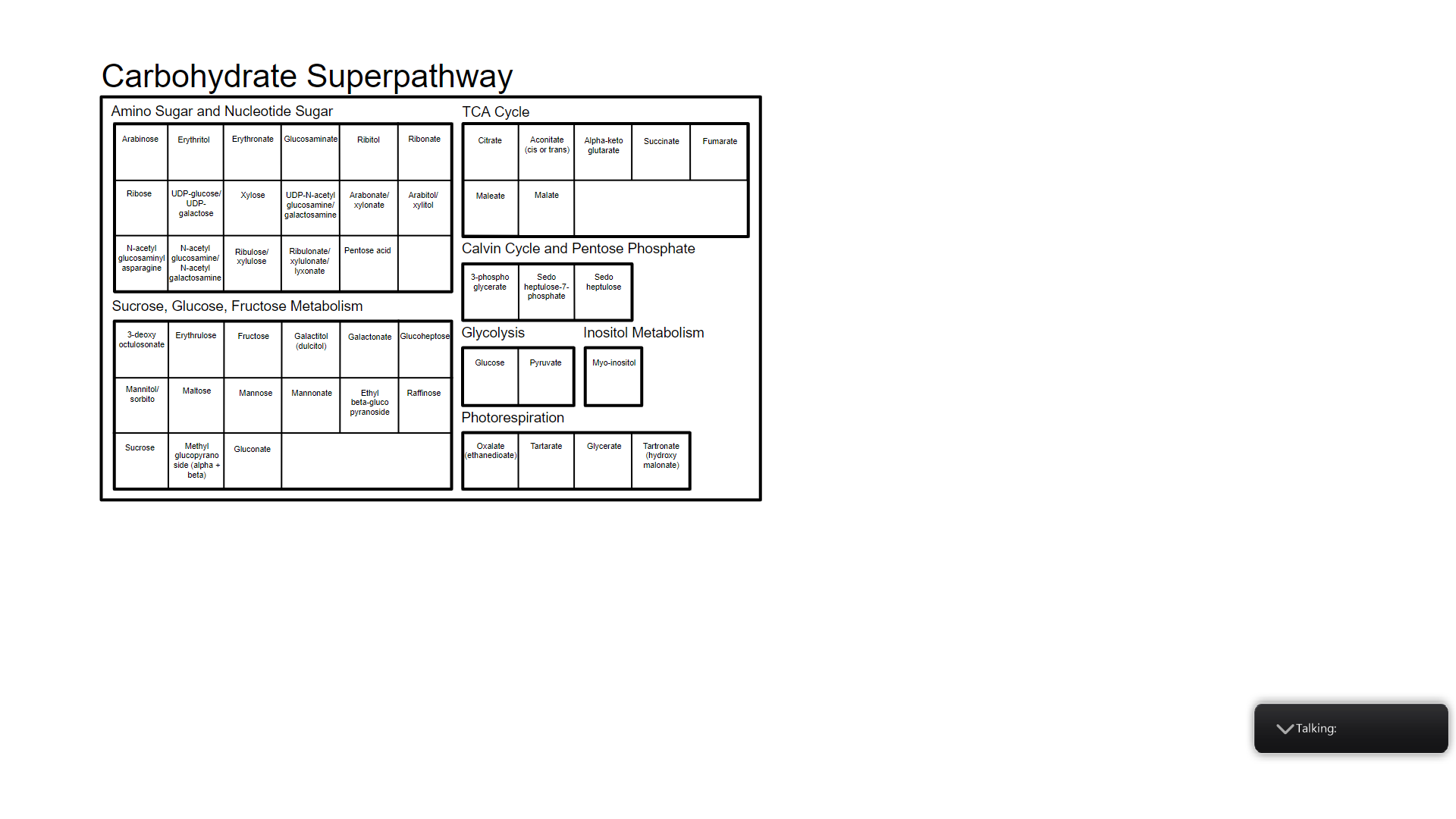

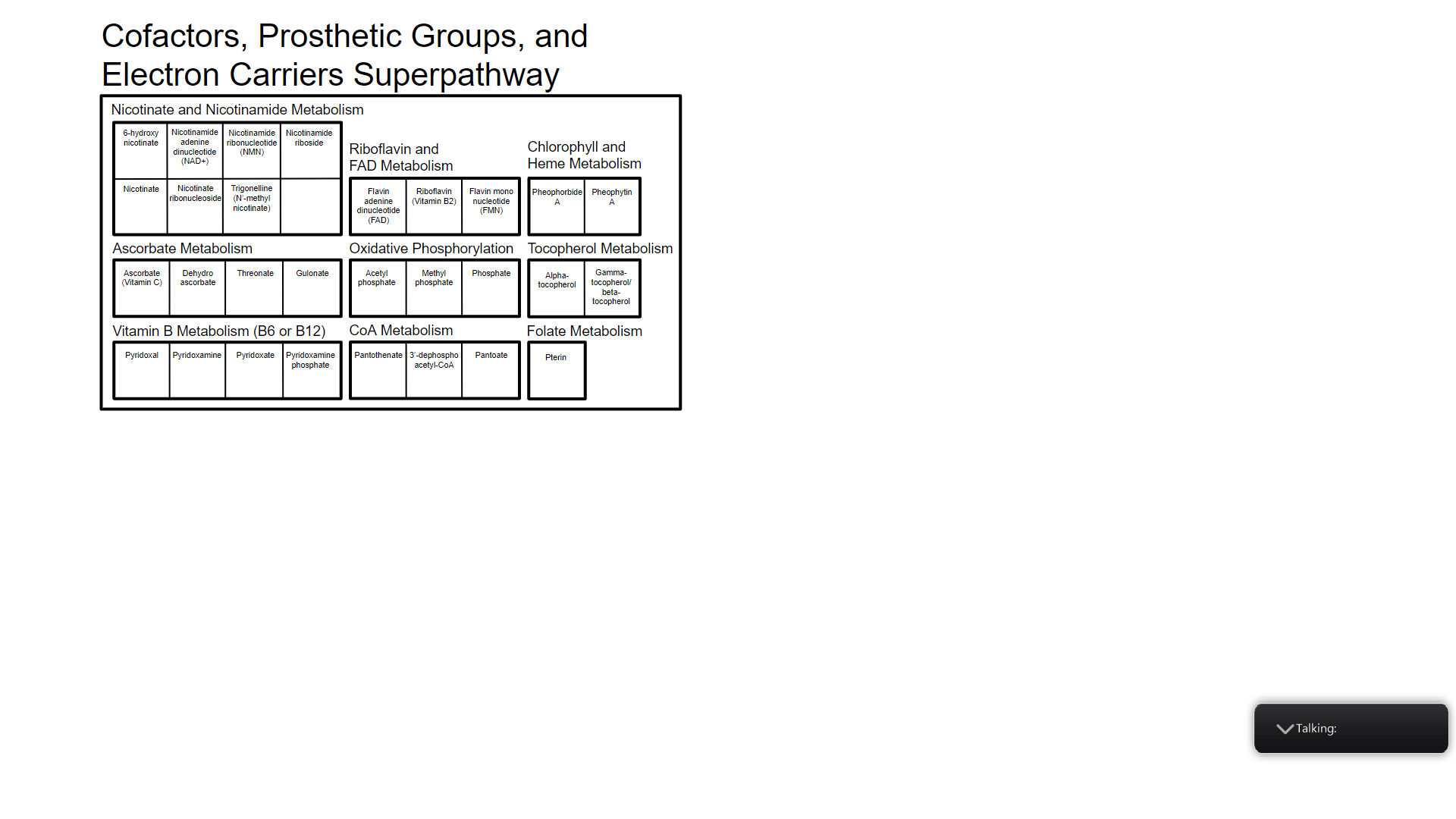

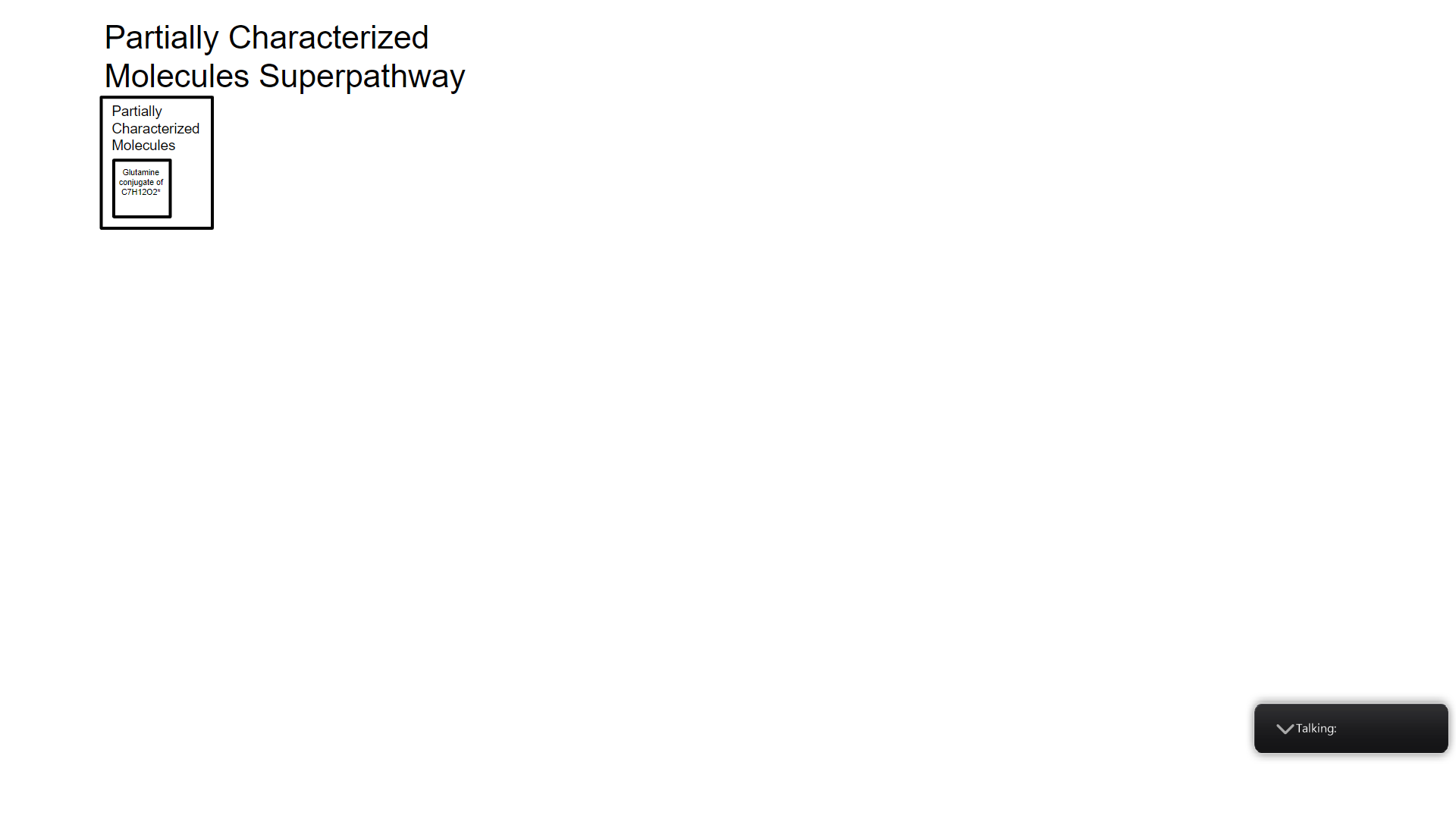

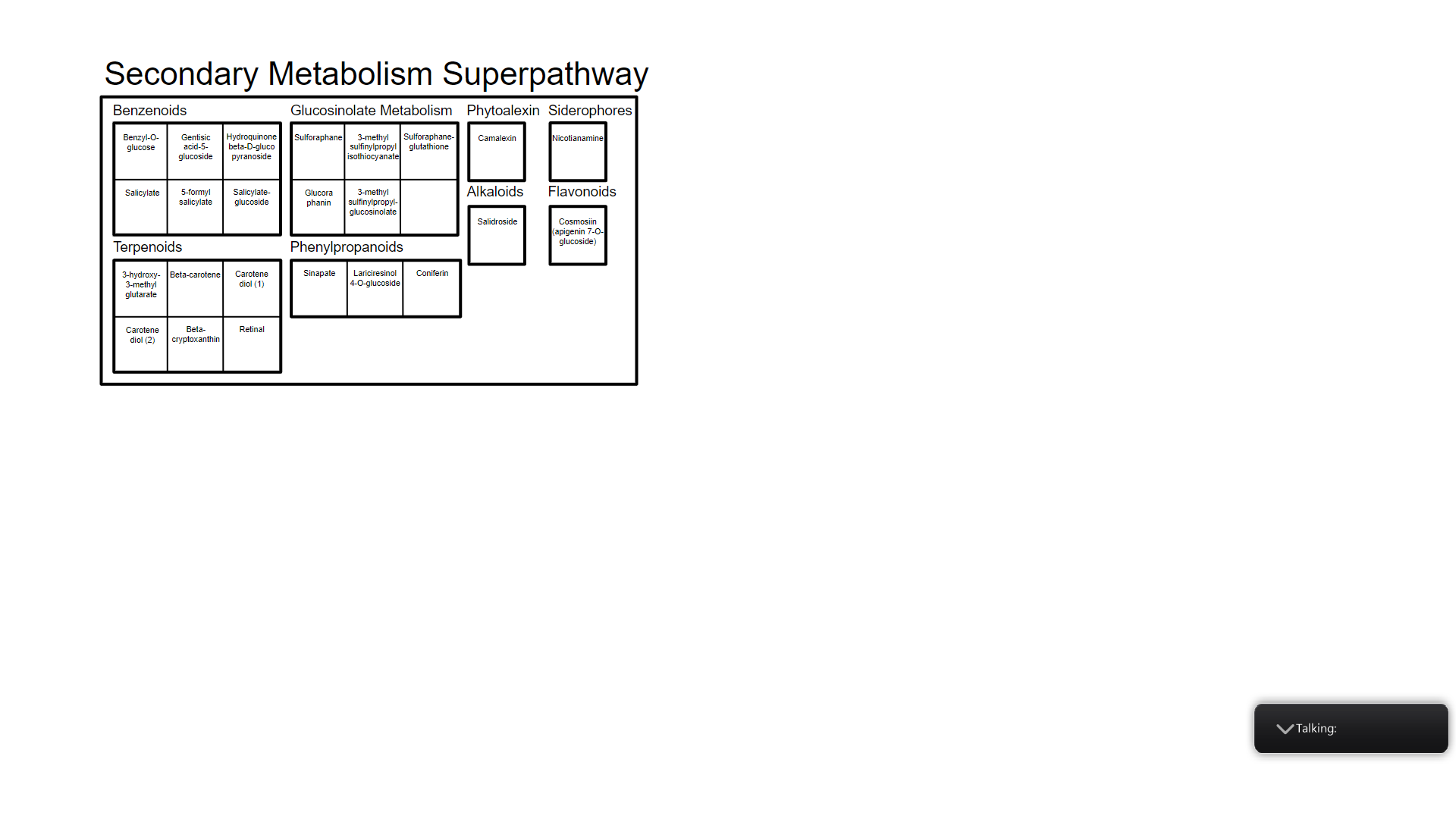

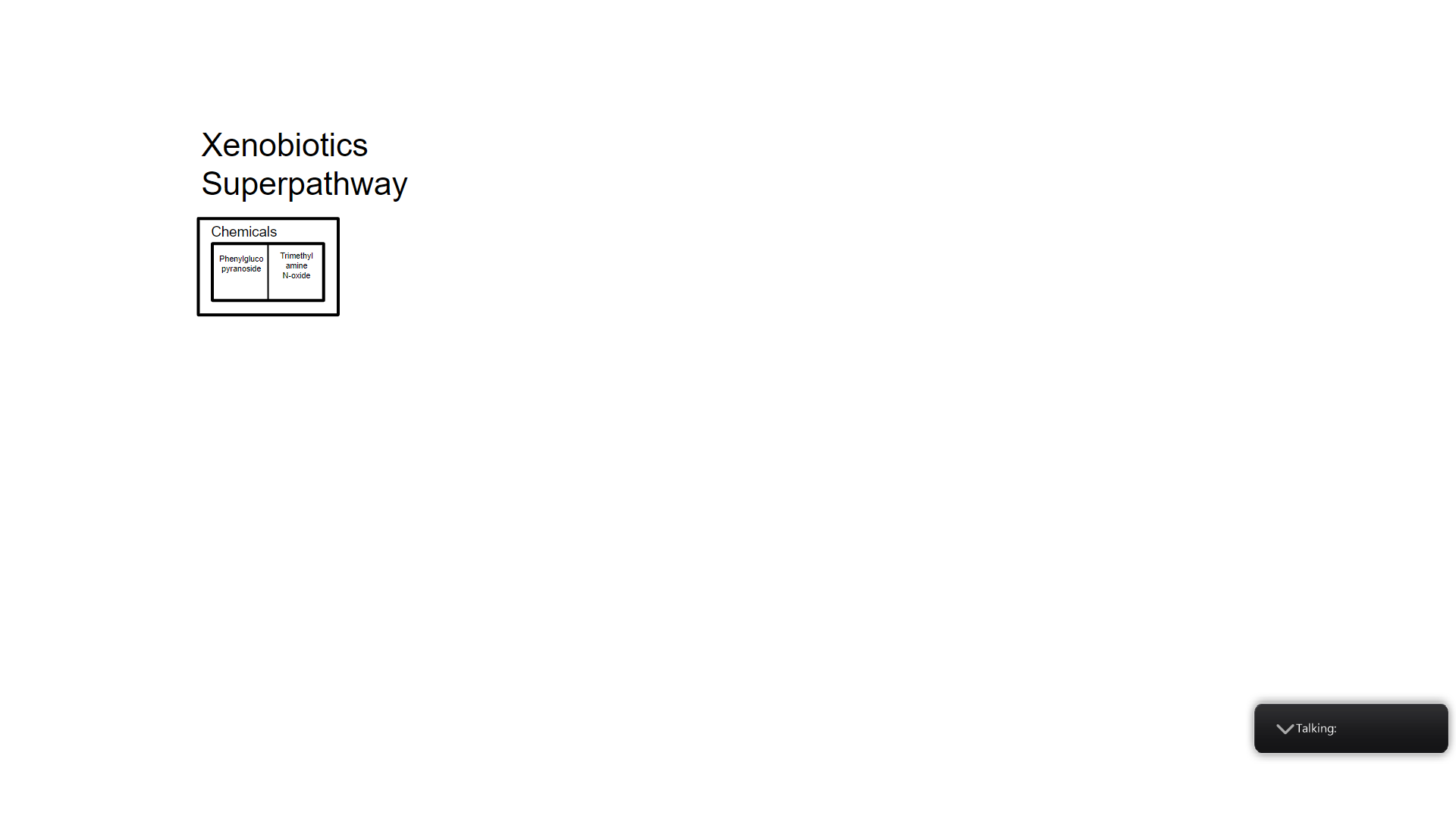

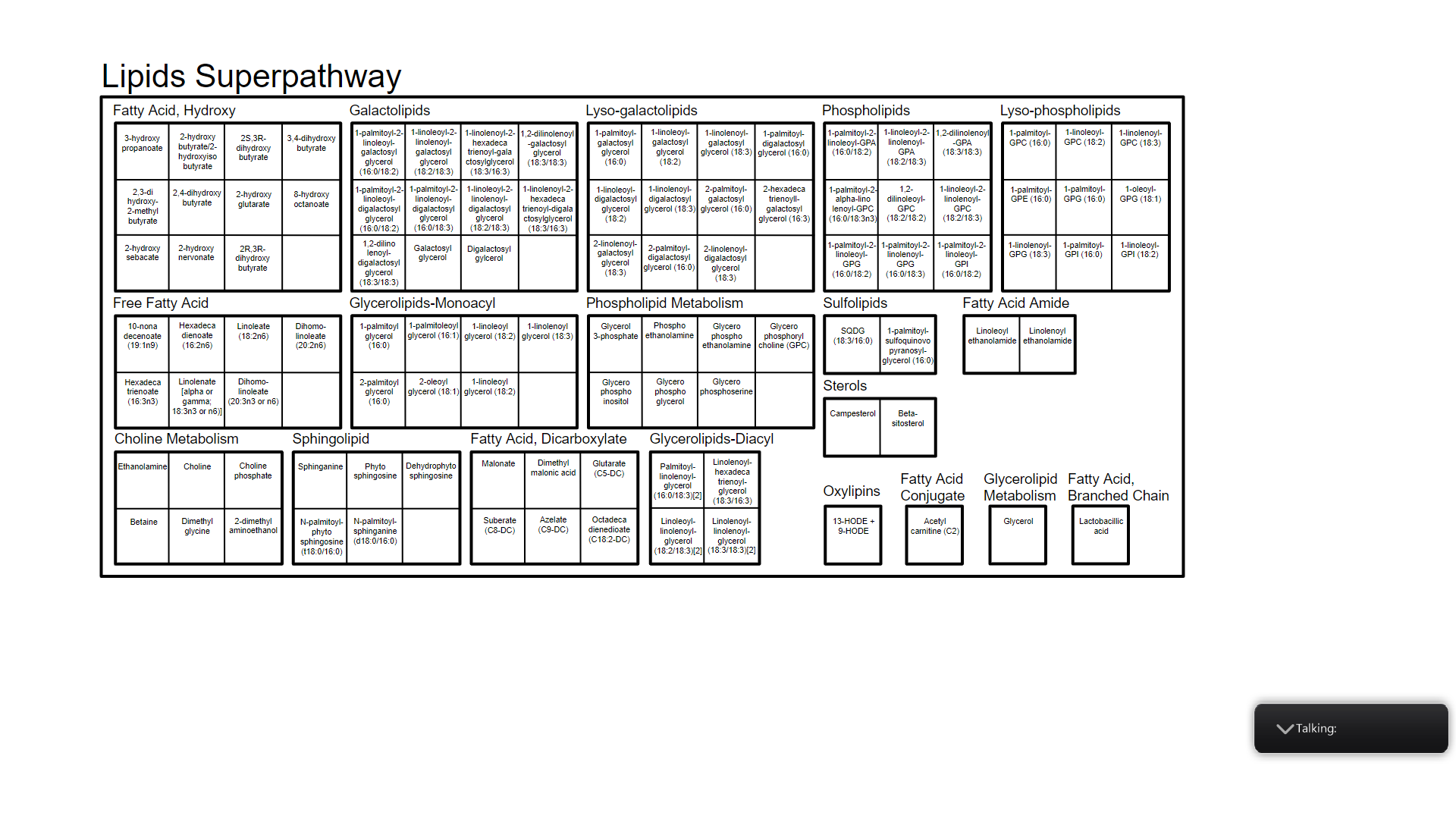

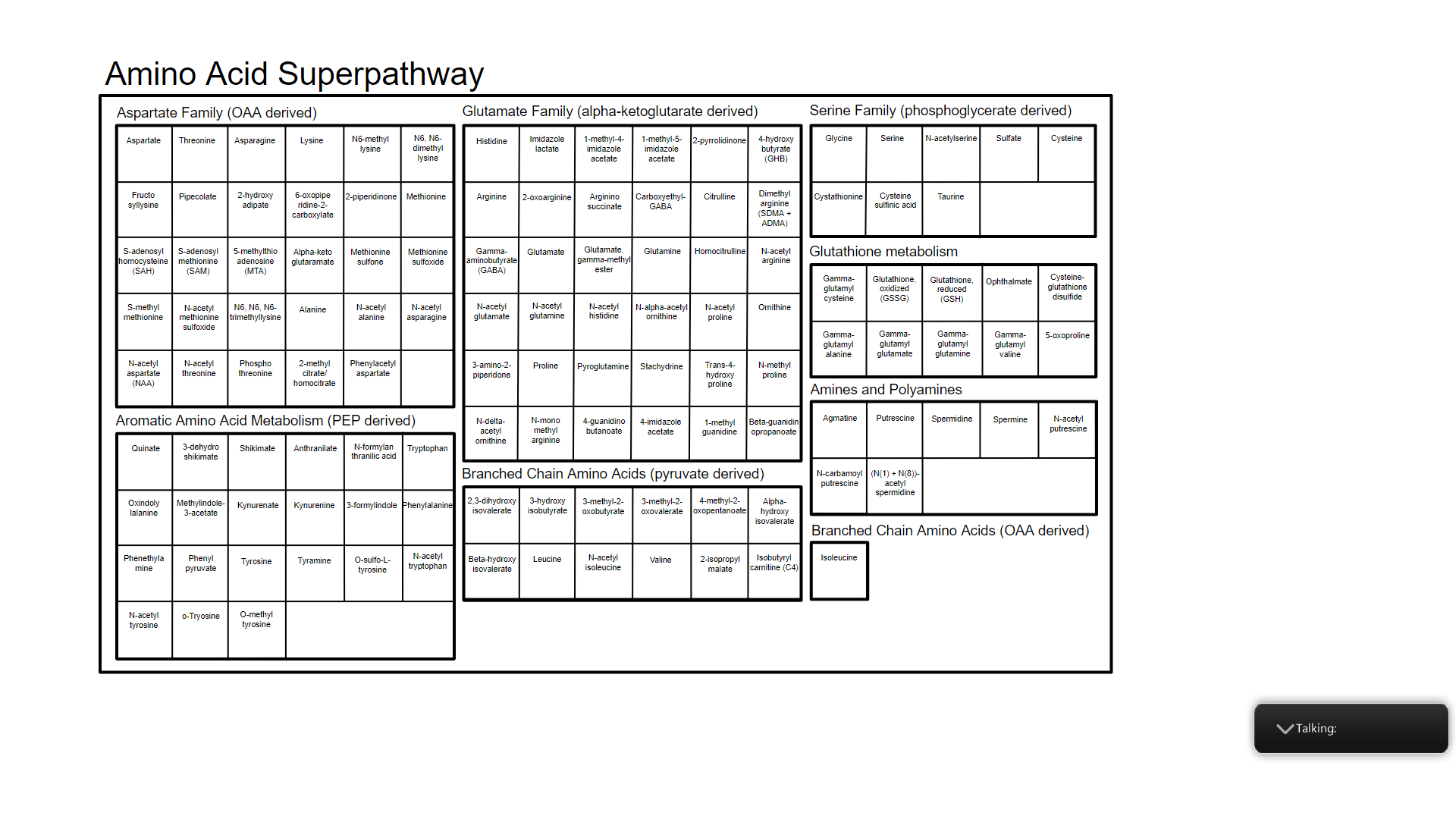

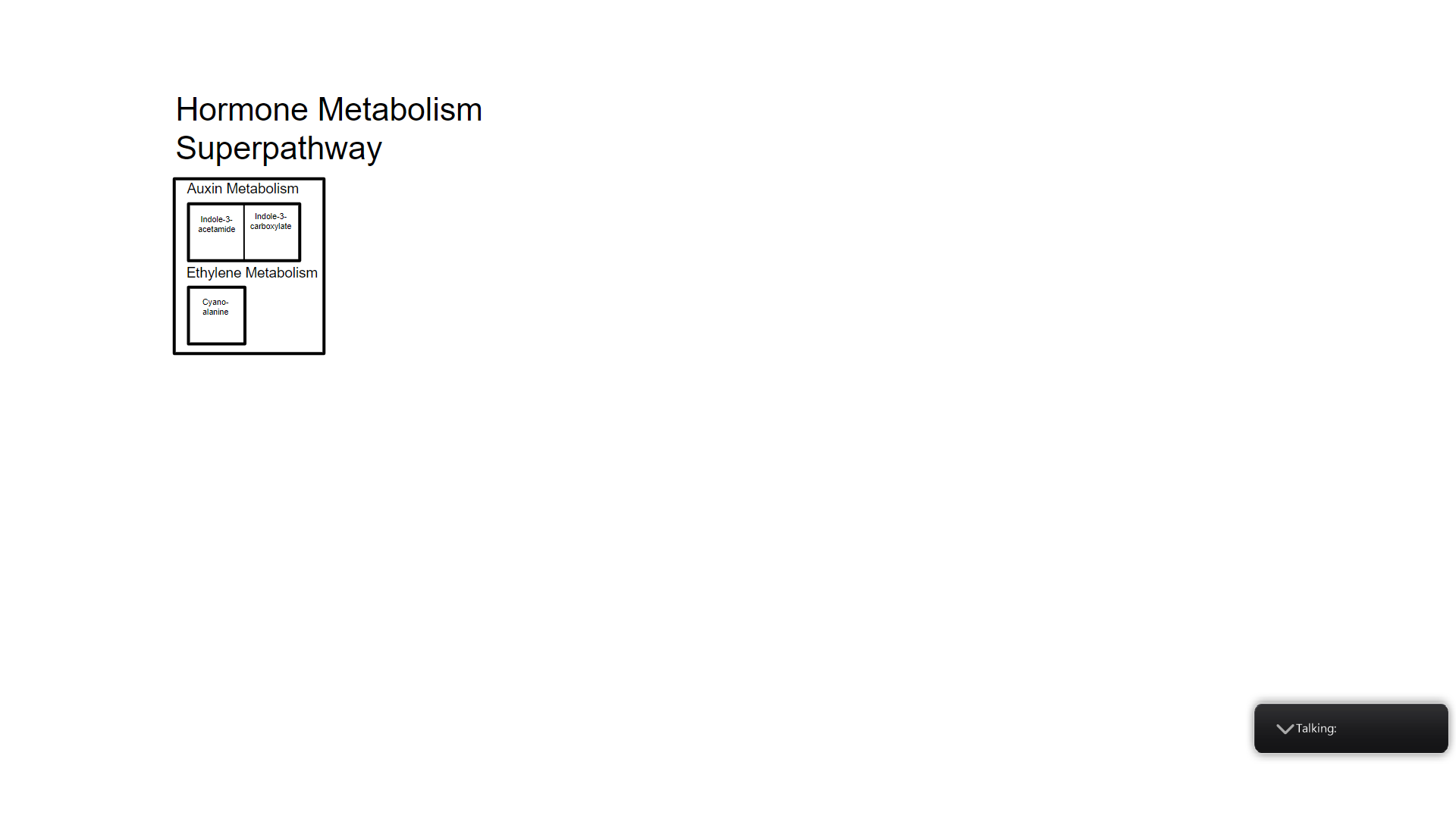

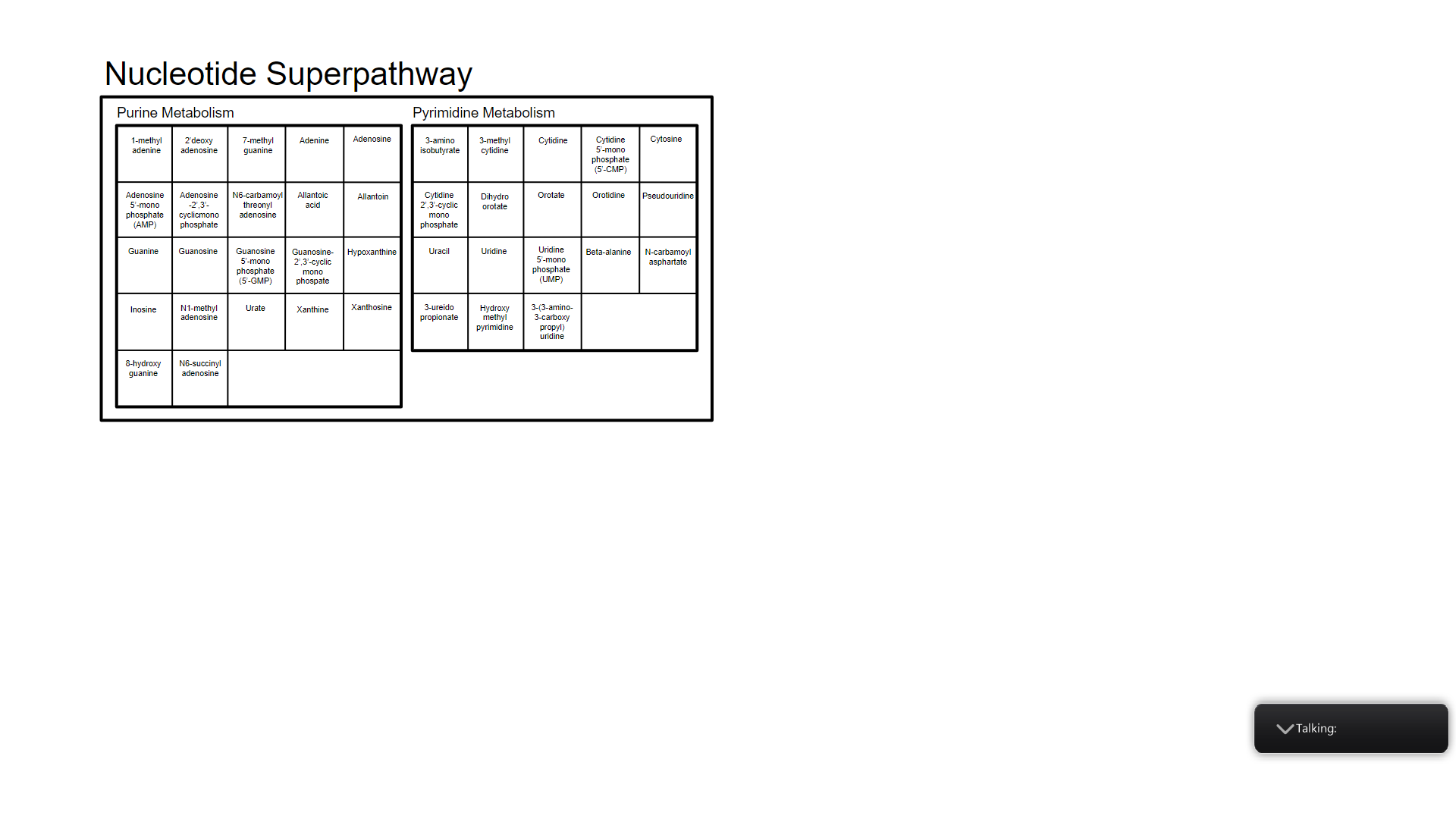

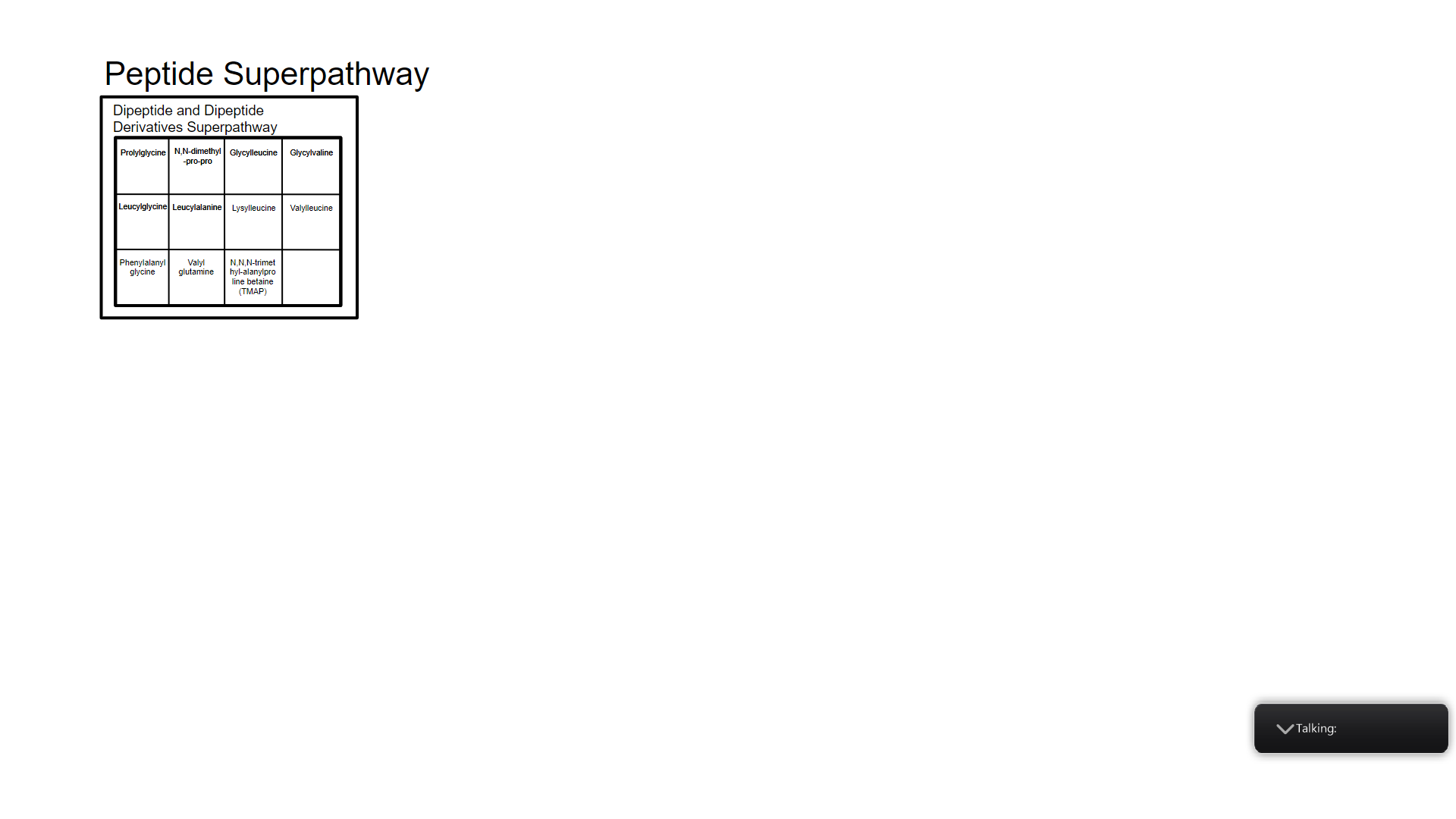

Supplementary Figure 2: Key to interpret the metabolic pathway groupings depicted in Figure 3 in the main text. Each metabolite name is written in each small square and each sub- and superpathway name is denoted outside of the boxes.
